# Survival and development of potato psyllid (Hemiptera: Triozidae) on Convolvulaceae: effects of a plant-fungus symbiosis (*Periglandula*)

**DOI:** 10.1101/371823

**Authors:** N. Kaur, W.R. Cooper, J.M. Duringer, I.E. Badillo-Vargas, G. Esparza-Díaz, A. Rashed, D.R. Horton

## Abstract

Plant species in the family Solanaceae are the usual hosts of potato psyllid, *Bactericera cockerelli* (Šulc) (Hemiptera: Psylloidea: Triozidae). However, the psyllid has also been shown to develop on some species of Convolvulaceae (bindweeds and morning glories). Developmental success on Convolvulaceae is surprising given the rarity of psyllid species worldwide associated with this plant family. We assayed 14 species of Convolvulaceae across four genera (*Convolvulus*, *Calystegia*, *Ipomoea*, *Turbina*) to identify species that allow development of potato psyllid. Two populations of psyllids were assayed (Texas, Washington). The Texas population overlaps extensively with native Convolvulaceae, whereas Washington State is noticeably lacking in Convolvulaceae. Results of assays were overlain on a phylogenetic analysis of plant species to examine whether Convolvulaceae distantly related to the typical host (potato) were less likely to allow development than species of Convolvulaceae more closely related. Survival was independent of psyllid population and location of the plant species on our phylogenetic tree. We then examined whether presence of a fungal symbiont of Convolvulaceae (*Periglandula* spp.) affected psyllid survival. These fungi associate with Convolvulaceae and produce a class of mycotoxins (ergot alkaloids) that may confer protection against plant-feeding arthropods. *Periglandula* was found in 11 of our 14 species, including in two genera (*Convolvulus*, *Calystegia*) not previously known to host the symbiont. Of these 11 species, leaf tissues from five contained large quantities of two classes of ergot alkaloids (clavines, amides of lysergic acid) when evaluated by LC-MS/MS. All five species also harbored *Periglandula*. No ergot alkaloids were detected in species free of the fungal symbiont. Potato psyllid rapidly died on species found to harbor *Periglandula* and fungus-produced alkaloids, but survived on species in which the mutualism was absent. These results support the hypothesis that a plant-fungus symbiotic relationship affects the suitability of certain Convolvulaceae to potato psyllid.

## Introduction

The potato psyllid, *Bactericera cockerelli* (Šulc) (Hemiptera: Psylloidea: Triozidae) is a pest of solanaceous crops such as potatoes, tomatoes, and peppers. The psyllid occurs throughout the western and central United States, Canada, Mexico, and Central America [1], and as an introduction in New Zealand and Australia [2, 3]. High densities of the psyllid may lead to plant disorders known as “psyllid yellows” [4, 5] caused by a toxin that is injected into plants during the psyllid’s feeding activities [6]. However, recent crop losses have been caused primarily by a bacterial pathogen, ‘*Candidatus* Liberibacter solanacearum’ (Lso), that is transmitted by the psyllid [1]. Difficulties in managing potato psyllid and its associated Liberibacter are in part due to poor understanding of the role that non-crop species have in the biology of the vector. Most species of psyllids are monophagous or oligophagous, limited to development on plants within a single genus or family [7, 8]. Potato psyllid is unusual in being able to develop on plants across more than a single family [9, 10, 11, 12]. Non-crop plant species act as reservoirs of the insect during the growing season and may help the psyllid bridge intervals in which crop hosts are unavailable [10, 13, 14, 15, 16, 17]. It is therefore important to know what non-crop species of plants found in potato or tomato growing regions also support the reproduction and development of potato psyllid.

Although plant species in the family Solanaceae (Solanales) are the typical developmental hosts for potato psyllid, at least some species in the Convolvulaceae (Solanales) also support development [9, 10, 11, 12, 14, 18]. Observations leading to this conclusion include rearing trials [9, 10, 11, 12] and field records [14, 18]. Developmental success on Convolvulaceae is unexpected given that Convolvulaceae is substantially underrepresented among plant families as hosts of Psylloidea. Despite its extensive diversity and widespread distribution [19] Convolvulaceae is listed as a developmental host for only five species of psyllids worldwide, including potato psyllid [20]. Rearing trials with potato psyllid have been limited to two species, *Convolvulus arvensis* L. (field bindweed) and *Ipomoea batatas* (L.) Lam. (sweet potato). While potato psyllid is able to complete development on these species, development rates are slow and may be accompanied by nymphal mortality [10, 12].

In this study, we examined the development of potato psyllid on species and genera of Convolvulaceae that have not previously been assayed. Our assays targeted species that are native to North America and are thus likely to have an evolutionary history with at least some populations of potato psyllid. Our first objective was to assay a taxonomically broader group of Convolvulaceae than previously done, to determine whether plant suitability extends beyond *C. arvensis* and *I. batatas*. Part of this objective included a comparison of two haplotypes of the psyllid on each plant species. Potato psyllid occurs as a minimum of four unique genetic types or “haplotypes” [21, 22] that we now know differ biologically [23, 24, 25, 26, 27]. We compared developmental success on Convolvulaceae between two of these haplotypes, the Central haplotype and the Northwestern haplotype. Convolvulaceae is highly diverse in the southern US and Mexico [28] where its presence overlaps extensively with the distribution of the Central haplotype [29]. In contrast, native Convolvulaceae are almost completely absent from the Pacific Northwest region of the US [28] where the Northwestern haplotype of potato psyllid is endemic [21, 22, 30]. Thus, the Northwestern haplotype is likely to have a much-reduced field history with native Convolvulaceae in comparison to the Central haplotype.

Our second objective was to look for traits that predict whether a given plant species allows psyllid development. We addressed two separate questions in this objective. First, we examined whether suitability is predicted by the location of plant species within a phylogenetic tree. Because of the strong tendency towards host specificity among species of Psylloidea, host switching or dietary expansion by psyllids tends to be phylogenetically conserved [31] such that evolutionary shifts in diets by psyllids are often between closely related plant species [8, 31, 32]. This specialism prompted us to examine whether plant suitability tracked plant phylogeny. We constructed a phylogenetic tree from DNA-sequence data to examine whether plant species allowing successful development of potato psyllid clustered together in the tree, as would be expected if plant chemistry or other traits affecting psyllid host use also grouped phylogenetically [33].

We then examined whether psyllid development was affected by the presence of a plant-fungus mutualism found in Convolvulaceae. The Convolvulaceae is unusual among dicotyledonous plant families in its association with a class of chemicals known as ergot alkaloids [34]. Many species of Convolvulaceae have formed a symbiotic association with clavicipitaceous fungi in the genus *Periglandula* [35, 36, 37, 38]. This fungus is vertically transmitted, and is present systemically in members of the family Convolvulaceae [39] often forming epiphytic colonies surrounding peltate glandular trichomes on the adaxial leaf surfaces [35, 40] where the colonies produce ergot alkaloids [41]. This symbiosis appears to be most common in *Ipomoea* and related genera, with possibly 450 or more plant species worldwide having the association [34, 42]. Similar alkaloids produced by clavicipitaceous fungi in grasses have been shown to have deleterious effects against herbivorous insects [43, 44, 45]. The defensive properties of ergot alkaloids associated with the Convolvulaceae-*Periglandula* symbiosis have received almost no attention, although extracts from *Ipomoea parasitica* (H.B.K.) G. Don, have been found to reduce feeding and digestive efficiency of caterpillars [46]. Our overall goal therefore was to examine whether survival and development of the potato psyllid was correlated with the presence or absence of *Periglandula*, and to determine whether psyllid development was affected by the types and quantities of fungal alkaloids produced by this symbiosis.

## Materials and Methods

### Source of plants and insects

Insect bioassays included a screening of 11 species of native Convolvulaceae distributed across three plant genera (*Convolvulus*, *Ipomoea*, and *Turbina*), and three introduced species in *Convolvulus* and *Calystegia* including the widespread pest field bindweed, *C. arvensis* (Table 1). All species except *Calystegia silvatica* overlap geographically with psyllids of the Central haplotype (Table 1). The Northwestern haplotype overlaps geographically with *C. arvensis*, possibly with *C. silvatica*, and is likely to have some overlap with the four species of *Ipomoea* that are grown extensively as summer ornamentals (Table 1). It is unlikely that these ornamentals are able to survive the winter conditions of the Pacific Northwest.

**Table 1.** List of plant species used in assays (origin, geographic overlap with psyllid haplotype, and source).

| Species | North America Origin | Overlap of insect haplotype and plant* | Source |
| --- | --- | --- | --- |
| *Convolvulus equitans* Benth. | Native | C | Western Region Plant Introduction Station, Pullman WA |
| *Convolvulus tricolor* L. ‡ | Introduced | C+N | J.L. Hudson, Seedsman, La Honda, CA |
| *Convolvulus arvensis* L. | Introduced | C+N | Prosser, WA |
| *Calystegia silvatica* (Kit.) Griseb.** | Introduced | N? | Tillamook Co., OR |
| *Ipomoea alba* L. ‡ | Native | C+N | The Sample Seed Shop, Buffalo, NY |
| *Ipomoea cordatotriloba* Dennstedt | Native | C | Georgia Vines, Claxton, GA |
| *Ipomoea hederacea* L. | Native | C | J.L. Hudson, Seedsman, La Honda, CA |
| *Ipomoea ternifolia* Torrey | Native | C | Southwest Seeds, Dolores, CO |
| *Ipomoea nil* (L.) Roth† | Native | C+N | The Sample Seed Shop, Buffalo, NY |
| *Ipomoea imperati* (Vahl) Grisebach | Native | C | South Padre Island, TX |
| *Ipomoea leptophylla* Torrey | Native | C | Georgia Vines, Claxton, GA |
| *Ipomoea pandurata* (L.) G.F. Meyer | Native | C | Georgia Vines, Claxton, GA |
| *Ipomoea tricolor* Cavanilles† | Native | C+N | J.L. Hudson, Seedsman, La Honda, CA |
| *Turbina corymbosa* (L.) Rafinesque | Native | C | J.L. Hudson, Seedsman, La Honda, CA |
| *Solanum tuberosum* L. | Native | C+N | Skone & Connors, Warden, WA |

\*C: Central haplotype occurs within geographic range of plant; N: Northwestern haplotype occurs within geographic range of plant. Haplotype distribution data [21, 22, 29, 30]. Plant distribution data [28].
\*\**Calystegia* is a taxonomically difficult genus with species often exhibiting substantial geographic variation in morphological traits [47]. We believe that the *Calystegia* assayed in this study is *Calystegia silvatica* (Kit.) Griseb. subsp. *disjuncta* Brummitt [48, 49]. *Calystegia silvatica* subsp. *disjuncta* is likely of Mediterranean origin, although there have been suggestions (probably incorrect) that it is native to North America [49]. It is unclear whether *C. silvatica* overlaps geographically with potato psyllid given historical uncertainties in distribution of the plant in the western U.S. ‡ In the Pacific Northwest plant is grown only as a summer ornamental. This may result in some level of sympatry with the Northwestern haplotype.

Test plants were examined in side-by-side comparisons with potato, *Solanum tuberosum* L. (‘Russet Burbank’) (Solanaceae), a typical and highly suitable host for potato psyllid. Plants were grown either from seeds or from stem cuttings (sources listed in Table 1). Seeds were scarified using sandpaper and soaked in gibberellic acid (1000 ppm in water) for 24 h prior to planting. Plants were grown in 10-cm pots (volume ~ 473.3 cm^3^) containing four parts commercial potting soil (Miracle-Gro Moisture Control Potting Mix, Scotts Company, Marysville, OH), one part perlite (Miracle-Gro Perlite, Scotts Company, Marysville, OH), and one part clean sand, and maintained in a greenhouse under ambient light supplemented with grow lights. Assays were done at the USDA –ARS in Wapato, WA. Plants at 1 to 4 fully expanded leaf stage were used in the assays.

Potato psyllids to be used in assays were obtained from colonies maintained at the USDA-ARS facility in Wapato, WA. The parental insects for colonies were collected from potato fields near Weslaco, TX in March 2017 (Central haplotype, APHIS permit P526P-17-00366) and from solanaceous weeds growing near Prosser, WA in the summer and autumn of 2016 (Northwestern haplotype). The colonies were maintained on potato (‘Russet Burbank’) at 22°C and a 16:8 h light: dark cycle. Colonies were assayed preceding the study using high resolution melting analysis to confirm haplotype status [21]. Colonies were checked periodically for Lso infection using PCR detection methods [50], and Lso free psyllids were used in these assays.

### Suitability of Convolvulaceae to potato psyllid

Our primary objective was to determine whether the potato psyllid is able to complete development on targeted plant species. Given the large size of the experimental design (two psyllid haplotypes x 15 plant species), we limited our measures of psyllid performance to two traits: egg-to-adult survival (as a yes/no variable), and egg-to-adult development time (in days). Ten adults (unsexed) from both haplotypes were collected from their respective colony cages. The ten psyllids of a given haplotype were confined for egg-laying on a test plant kept individually in a 7.5 L plastic container (Cambro^®^, Huntington Beach, CA) modified to allow ventilation at 22°C and a 16:8 h light: dark cycle. Once 20 or more eggs were present on a test plant, the adults were removed. Containers were monitored every 2-3 days for hatching of eggs and subsequent development of nymphs.

For plant species on which psyllids developed successfully, we recorded the number of days required to develop from egg deposition to production of the first adult (i.e., the minimum time required to complete development). Once new adults were seen in a container, that plant and container was dismantled. A leaf was collected from the plant for DNA extraction and biochemical analysis (described below).

On species which failed to support development, mortality almost invariably occurred as first instar nymphs often within 48 h of hatch. When monitoring showed that all nymphs on a given plant were dead, the assay for that plant and container was dismantled, and leaf samples were collected for DNA extraction and ergot alkaloid quantification. We had five replicates per plant species per psyllid haplotype combination. The large number of treatments, combined with uneven germination of seed, did not allow us to conduct the five replicates simultaneously. Thus, each replicate was initiated on a separate date, with date of assay included in the statistical analyses as a blocking factor (see Statistical analyses).

### Phylogenetic mapping of Convolvulaceae

DNA was extracted using a cetyltrimethylammonium bromide (CTAB) precipitation method [51]. Two different universal plant barcoding primer sets were used. The first primer set targeted approximately 500 bp of the internal transcribed spacer region (*ITS*): ITS2F (ATGCGA TACTTGGTGTGAAT) and ITS3R (GACGCTTCTCCAGACTACAAT) [52]. The second primer set targeted approximately 684 bp region of the chloroplast maturase K gene (*matK*): matK 472-F (CCCRTYCATCTGGAAATCTTGGTT) and matK 1248-R (GCTRTRATAATGAGAAAGATTTCTGC) [53]. PCR conditions used for both primer sets were similar, consisting of an initial denaturation step of 94°C for 5 min followed by 35 cycles of 94°C for 30 s, 56°C for 30 s, and 72°C for 42 s, followed by a final extension at 72°C for 10 min. Each 20μl reaction contained Amplitaq Gold 360 PCR Master Mix (Invitrogen, Carsbad, CA), 500nM of each primer, and DNA template (10-20 ng). Upon amplification, bands were excised from agarose gels, purified using GenElute minus ethidium bromide spin columns (Sigma, St. Louis, MO), and were cloned using a TOPO TA cloning kit with TOP10 *E. coli* chemical competent cells (Invitrogen, Carlsbad, CA). The QIAprep spin mini prep kit (Qiagen, Valencia, CA) was used to prepare plasmid DNA for sequencing by MC Laboratories (MC Lab, San Francisco, CA). Sequences were deposited into GenBank (Table 2).

**Table 2.** GenBank Accession Numbers

| Species | ITS | matK | dmaW |
| --- | --- | --- | --- |
| *Convolvulus equitans* | MG889580 | MH198126 | MH195190 |
| *Convolvulus tricolor* | MG889582 | MH198117 | MH195191 |
| *Convolvulus arvensis* | MG889579 | MH198115 | nd |
| *Calystegia silvatica* | MG889581 | MH198116 | MH195189 |
| *Ipomoea alba* | MG910322 | MH198118 | nd |
| *Ipomoea cordatotriloba* | MG910323 | MH198119 | MH195192 |
| *Ipomoea hederacea* | MG910324 | MH198127 | MH195193 |
| *Ipomoea ternifolia* | MG910327 | MH198121 | MH195196 |
| *Ipomoea nil* | MG910328 | MH198122 | nd |
| *Ipomoea imperati* | MG910325 | MH198120 | MH195194 |
| *Ipomoea leptophylla* | MG910326 | MH198128 | MH195199 |
| *Ipomoea pandurata* | MG910329 | MH198123 | MH195195 |
| *Ipomoea tricolor* | MG910330 | MH198124 | MH195197 |
| *Turbina corymbosa* | MG910332 | MH198125 | MH195198 |
| *Solanum tuberosum* | MG910331 | MH198129 | nd |

Not detected (nd)

DNA sequences were aligned and consensus sequences were made using Geneious R10 software (North America Biomatters Inc, Newark, NJ). The phylogenetic tree was constructed using a Tamura-Nei model and neighbor-Joining method with the Tree Builder function of Geneious R10 [54]. Phylogenetic distances for tree construction were estimated based upon concatenated sequences of *ITS* and *matK* regions. Potato was treated as an outgroup.

### Detection of the Convolvulaceae-*Periglandula* association

Because *Periglandula* is not always readily visible on plants even when the fungus is present, extraction and analysis of DNA-sequences is often used to confirm infestation. Presence or absence of the *dmaW* gene, encoding 4-(γ,γ –dimethylallyl) tryptophan synthase and required for the determinant step of ergot alkaloid synthesis, was evaluated using PCR [35, 55]. Plant DNA extracted using CTAB method was used to amplify approximately 1050 bp region using dmaWF5 (GACCGTAAACGAGTCAGGAA) and dmaWR2 (AAATACACCTGGGGCTCG) primers. PCR conditions consisted of an initial denaturation step of 95°C for 5 min followed by 40 cycles of 95°C for 1 min, 52°C for 1 min, and 72°C for 45 s, followed by a final extension at 72°C for 5 min. Each 20μl reaction contained Amplitaq Gold 360 PCR Master Mix (Invitrogen, Carsbad, CA), 500nM of each primer, and DNA template (10-20 ng). Upon amplification, bands were excised from agarose gels, purified using GenElute minus ethidium bromide spin columns, and were cloned (methods described in previously). Sequencing again was done by MC Laboratories. Sequences were deposited into GenBank (Table 2).

### Quantification of ergot alkaloids

Acetonitrile (ACN) and methanol (LC-MS grade) as well as acetic acid (OmniTrace Ultra), were purchased from EMD Millipore (Darmstadt, Germany). Ammonium acetate (>99.0%, HPLC grade) was obtained from Sigma Aldrich (St. Louis, MO USA). Ergot alkaloid standards were purchased from Romer Labs (Tulln, Austria) (biopure mix α-ergocornine, ergocristine, α-ergocryptine, ergometrine, ergosine and ergotamine) and Sigma-Aldrich (St. Louis, MO USA) (ergonovine, agroclavine, lysergic acid and lysergol). Ultrapure 18 mΩ cm-1 water was obtained from an Elga (Marlow, Buckinghamshire, UK.) PURELAB Ultra Genetic system.

Fully expanded leaves were collected from assayed plants at the end of suitability tests and subjected to air drying at the room temperature ~22-25°C for 3-5d. Dried tissue was ground using either a mortar and pestle or a cyclone sample mill with a 0.5 mm screen (UDY Corporation, Fort Collins CO). Extraction solution (79:20:1 ACN:water:acetic acid) was added to ground sample at a ratio of 4 mL/g and turned for 90 min in the dark [56]. The sample was then centrifuged for 2 min at 1462 x *g*. Dilution solution (250 μL 20:79:1 ACN:water:acetic acid) was added to 250 μL supernatant, vortexed for 10 sec, then placed in an amber HPLC vial for ergot alkaloid analysis by LC-MS/MS.

An ABI/SCIEX 3200 QTRAP LC-MS/MS system (Applied Biosystems, Foster City, CA USA) was used to monitor for ergoline and ergopeptide compounds via positive electrospray ionization, with separation performed using a Perkin Elmer (Waltham, MA USA) Series 200 autosampler and HPLC connected to a Gemini C18 column (150 x 4.6 mm, 5 μ, Phenomenex (Torrance, CA USA)) with a 4 x 3 mm security guard cartridge of similar packing [56]. Mobile phases consisted of 5 mM ammonium acetate and methanol:water:acetic acid in a ratio of 10:89:1(v/v/v) (A) or 97:2:1 (B) and were run in a gradient program at 1 mL/min. Multiple reaction monitoring (MRM) of two transitions (quantitative and qualitative) per compound was used to detect the ergot alkaloids ergonovine, ergotamine, ergocornine, α-ergocryptine, ergocristine, ergovaline, ergine, ergosine and their epimers, as well as agroclavine, chanoclavine, lysergol, lysergic acid, oxidized luol, dihydrolysergol, chanoclavine, dihydroergosine, dihydroergotamine, festuclavine, fumigaclavine and elymoclavine.

The presence of a mycotoxin was confirmed when the signal was equal to or greater than a signal-to-noise (S/N) ratio of 3:1 (limit of detection (LOD)), and both quantitative and qualitative transitions were present. Samples were quantitated blind as to sample identity against a standard curve using Analyst 1.6.2 and MultiQuant 3.0.1 (Applied Biosystems). The limit of quantitation (LOQ) was defined as the concentration at which the analyte had a precision and accuracy that did not exceed greater than 20% of the coefficient of variation [57]. LOD and LOQ for detected mycotoxins were as follows: ergotamine, ergocornine, ergosine ergocristine and agroclavine (1, 1 ng/mL); ergonovine (1, 2 ng/mL); lysergol (2, 2 ng/mL); lysergic acid (20 and 50 ng/mL). No commercial standards were available for chanoclavine, festuclavine, elymoclavine, elymoclavine fructoside, ergine and dihydrolysergol. These compounds were compared on a scale of present (“+” indicating low, “++” indicating high) or not present (“-“) amongst the plant species extracted based on relative peak area.

### Statistical analyses

Effects of plant species and psyllid haplotype on mean psyllid development time were examined using a generalized linear mixed model (Proc GLIMMIX) [58]. Plant species, psyllid haplotype, and the species x haplotype interaction were included as fixed effects, and replicate (N=5) was included as a random effect. The analysis was limited to plant species on which psyllids completed development to the adult stage. We specified an underlying gamma distribution using a DIST=gamma statement. This distribution is useful for modeling time-to-occurrence data [59]. The ILINK function was used to back-transform means into the original units (number of days to first adult). The CONTRAST statement was used to examine *a priori* defined comparisons among plant species following a significant plant species effect in the overall model (see results). The survival data (yes/no) were not analyzed statistically, as the two haplotypes showed identical results as to what plant species supported development (see results).

Genetic distances between species in a tree were calculated automatically by the Geneious R10 software using the Tamura-Nei model. This approach expresses distance as nucleotide substitutions per site. We used these distances to determine whether plant suitability for psyllids decreased as genetic distance from potato increased. We conducted a two-sample t-test [58] to determine whether mean genetic distance from potato differed between plant species allowing development and plant species not allowing development. If suitability was affected by genetic distance from the typical host (potato), we expected mean distance to be smaller for plants on which psyllids survived than plants on which psyllids failed to survive.

## Results

### Psyllid developmental success and plant phylogeny

Eggs were present within 24-48 h of adding egg-laying psyllids on all plant species except for *Ipomoea pandurata* and *Turbina corymbosa* (which required 72 h before eggs were present). Psyllids of both haplotypes failed to complete development in all five replicates on *Convolvulus equitans*, *Calystegia silvatica*, *Ipomoea imperati*, *Ipomoea leptophylla*, *Ipomoea pandurata*, *Ipomoea tricolor*, and *Turbina corymbosa*. Nymphs invariably died within a week of hatch on these species. On some species, notably *I. imperati*, *I. pandurata*, and *T. corymbosa*, mortality occurred within 24-48 h of egg hatch. Psyllids of both haplotypes completed development on potato, *Convolvulus arvensis*, *Convolvulus tricolor*, *Ipomoea alba*, *Ipomoea cordatotriloba*, *Ipomoea hederacea*, *Ipomoea ternifolia*, and *Ipomoea nil*.

A tree generated from *ITS* and *matK* sequences resolved the 14 species of Convolvulaceae into two major groups (Fig 1: Convolvuleae, Ipomoeeae), consistent with subfamilial groupings shown elsewhere in substantially more detailed taxonomic work [60]. The assay data were overlain on the tree to search for evidence that plant phylogeny predicted survival. Within Tribe Convolvuleae, psyllids of both haplotypes developed successfully on *C. arvensis* and *C. tricolor*, but failed to survive on *C. silvatica* and *C. equitans*, despite their phylogenetic closeness to *C. arvensis* (Fig 1). Within Tribe Ipomoeeae, psyllids of both haplotypes developed successfully on five species of *Ipomoea*, but failed to develop on four other *Ipomoea* or on *T. corymbosa*. Phylogenetic distance from the control host plant (potato) was calculated for each species of Convolvulaceae in the phylogenetic tree (distance matrix generated automatically by Geneious^®^, data not shown). A two-sample t-test demonstrated that mean phylogenetic distance from potato was statistically identical between plant species that allowed psyllid survival versus species on which psyllids failed to survive (P = 0.4563; Fig 2), confirming observations in the phylogenetic tree that phylogenetic nearness of a species to potato did not predict whether the psyllid would complete development on the plant (Fig 1).

**Fig 1.**
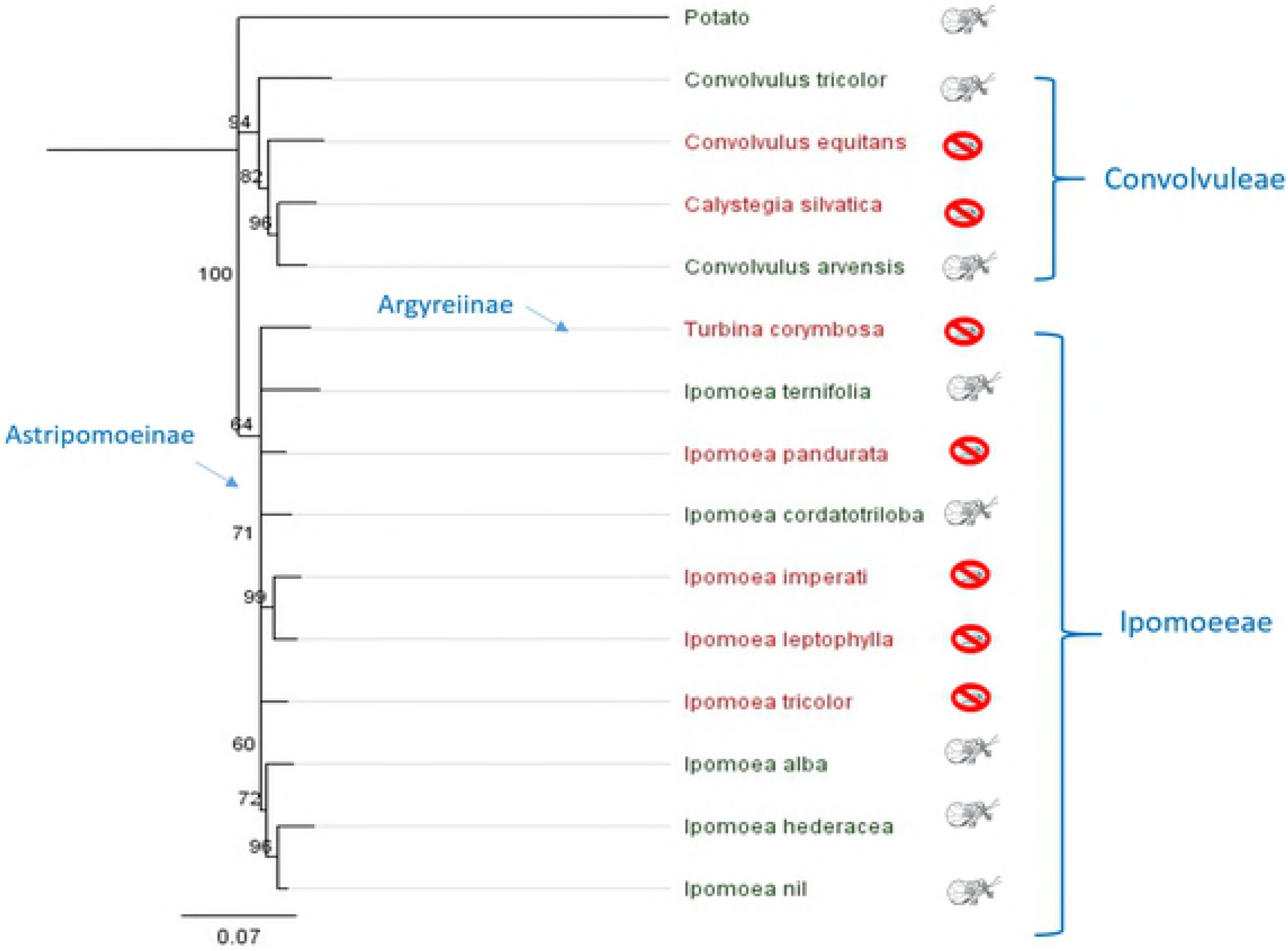
Phylogeny of assayed Convolvulaceae based on *ITS* and *matK* sequences. Node confidence was calculated using Neighbor Joining tree (Bootstrap replicates= 100). Species in red font followed by an insect kill icon failed to allow survival to adult stage; species in green font followed by a psyllid adult icon allowed egg-to-adult development.

**Fig 2.**
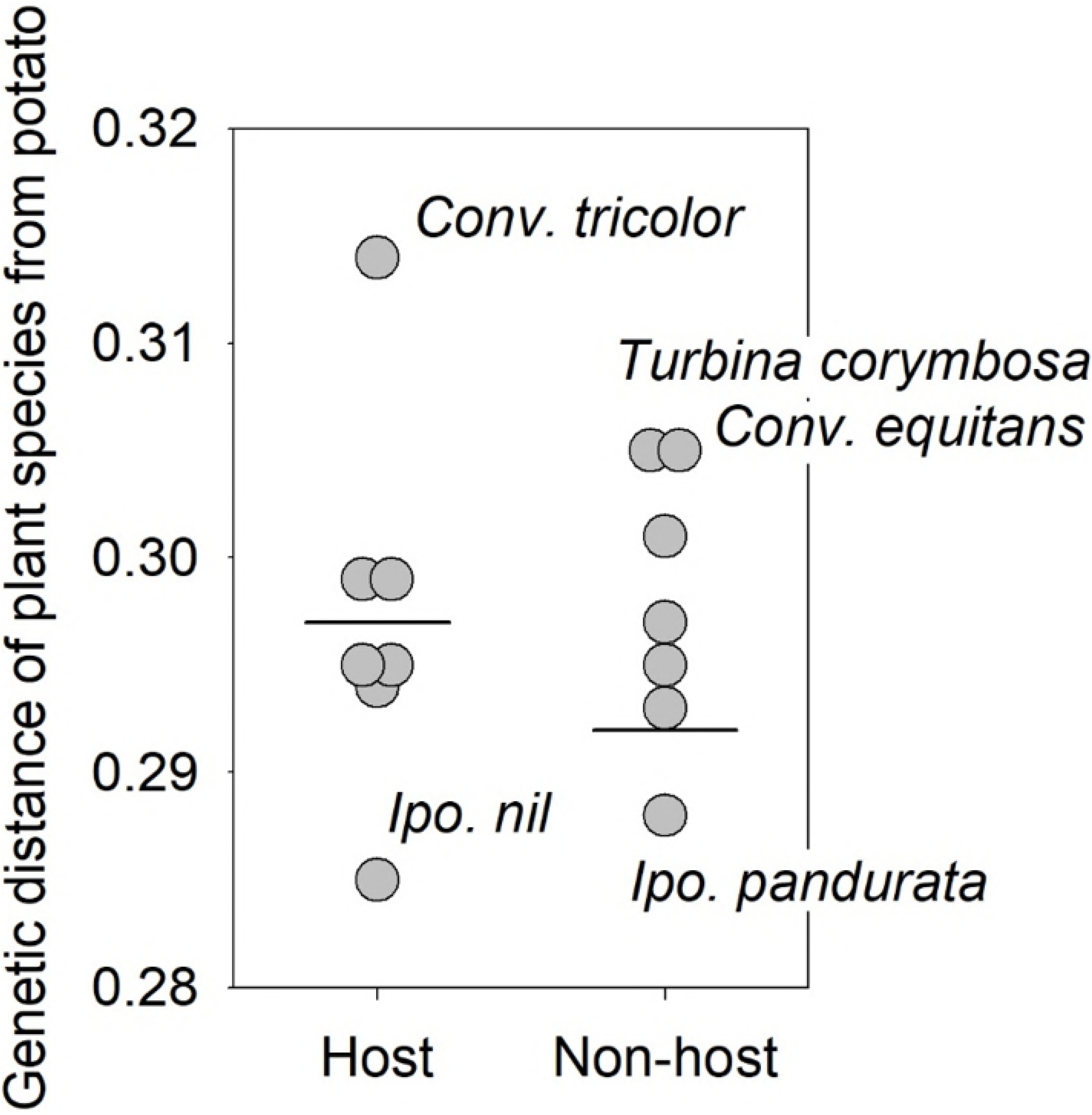
Scatter plot showing relationship between genetic distance of plant from potato (control) and survival of potato psyllid to the adult stage. Horizontal lines indicate mean distances.

We did not record actual rates of survival, so the assays cannot tell us whether percent survival on plant species that allowed egg-to-adult development was similar to survival on potato. To determine if developmental rates on Convolvulaceae were similar to rates on potato, we compared developmental times (number of days from oviposition to production of the first adult) of psyllids on potato and on those Convolvulaceae allowing survival to the adult stage. Development times varied between ~20-35 days depending upon psyllid haplotype and plant species (Fig 3). Mean development times differed statistically between psyllid haplotypes (*F* = 15.5; df = 1, 57.0; P <0.001) and among plant species (*F* = 2.8; df = 7, 57.1; *P* = 0.013); the haplotype x plant species interaction was not significant (*F* = 1.4; df = 7, 57.1; *P* = 0.22) indicating that the effects of plant species on psyllid development time was similar between the two haplotypes. The Central haplotype developed more rapidly (mean= 24.7 ± 1.1 d) than the Northwestern haplotype (mean = 29.4 ± 1.3 d), when averaged across host plant. We extracted contrasts to examine two *a priori* defined comparisons of interest. A test of mean development time on potato vs. Convolvulaceae was significant (*F* = 18.01; df = 1, 57.02; *P* < 0.001), and showed that mean development time on potato was statistically shorter than development time on Convolvulaceae (Fig 3). A second set of contrasts was extracted to examine whether there was evidence for plant effects within the Convolvulaceae, ignoring potato. Averaged over the two haplotypes, there was no evidence that development time of psyllids varied among species of Convolvulaceae (Fig 3: *F* = 0.31; df = 6, 57.1; *P* =0.93).

**Fig 3.**
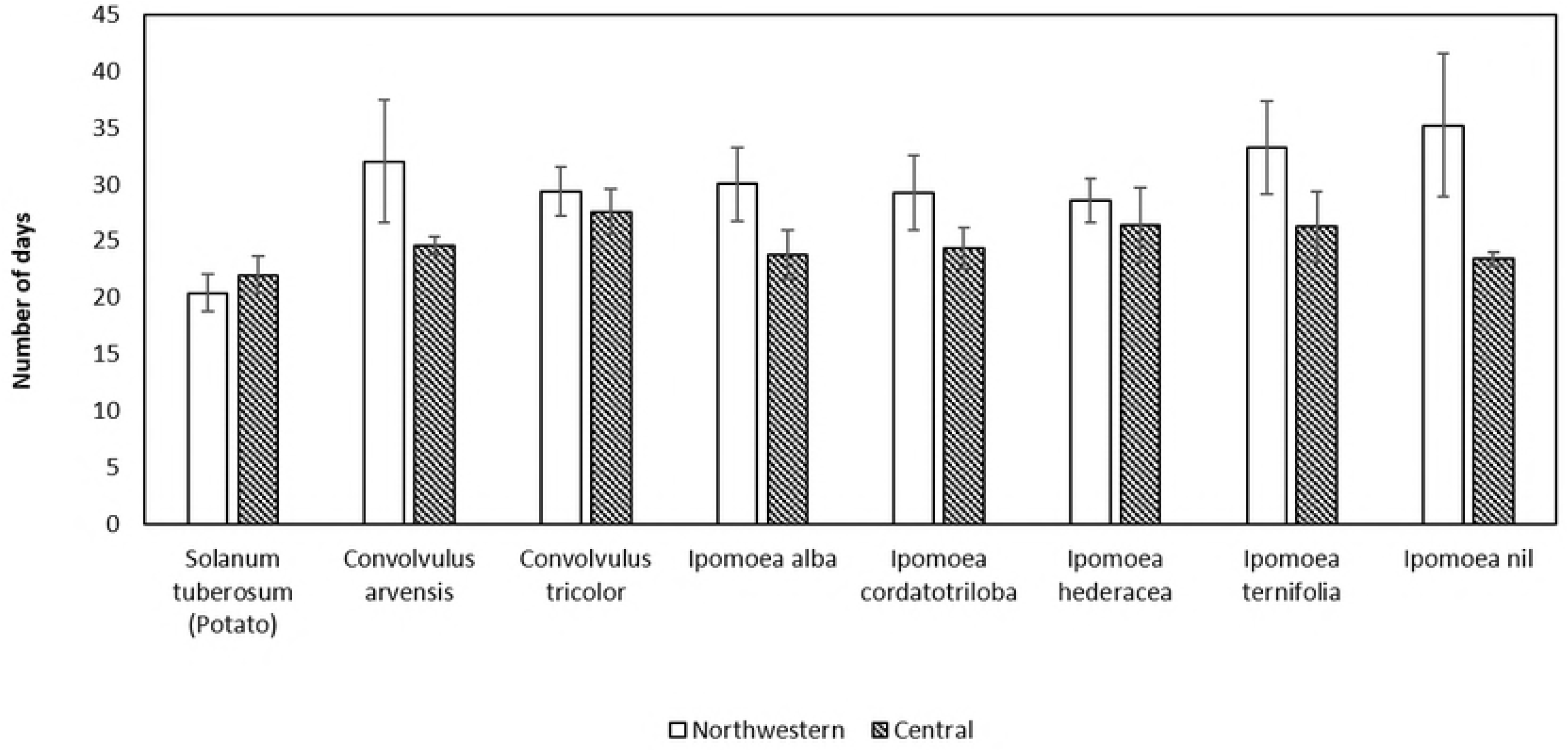
Number of days required to complete development from egg to adult stage by psyllids of the Northwestern and the Central haplotypes on potato and Convolvulaceae. Error bars represent standard error of mean.

### Psyllid developmental success and a plant-fungus symbiosis

Visible evidence for the presence of *Periglandula* was most pronounced in two species, *T. corymbosa* and *I. leptophylla* (Fig 4 AB). The fungal colonies were found on the adaxial surfaces of younger leaves. Because visible evidence for presence of fungal colonies was rare, we used a molecular approach for detection of the fungus. Analysis of DNA-sequences led to detection of the *dmaW* gene in 11 of 14 plant species (Fig 4 CD; Table 2), indicating widespread presence of *Periglandula* across species despite absence of visible evidence. Only three species (*C. arvensis*, *I. alba*, *I. nil*) failed to show presence of *Periglandula*. Presence of *Periglandula* in *Convolvulus* and *Calystegia* (Convolvuleae) was unexpected, as there had been no previous unambiguous evidence suggesting an association between *Periglandula* and plant species outside of the Ipomoeeae [34, 42].

**Fig 4.**
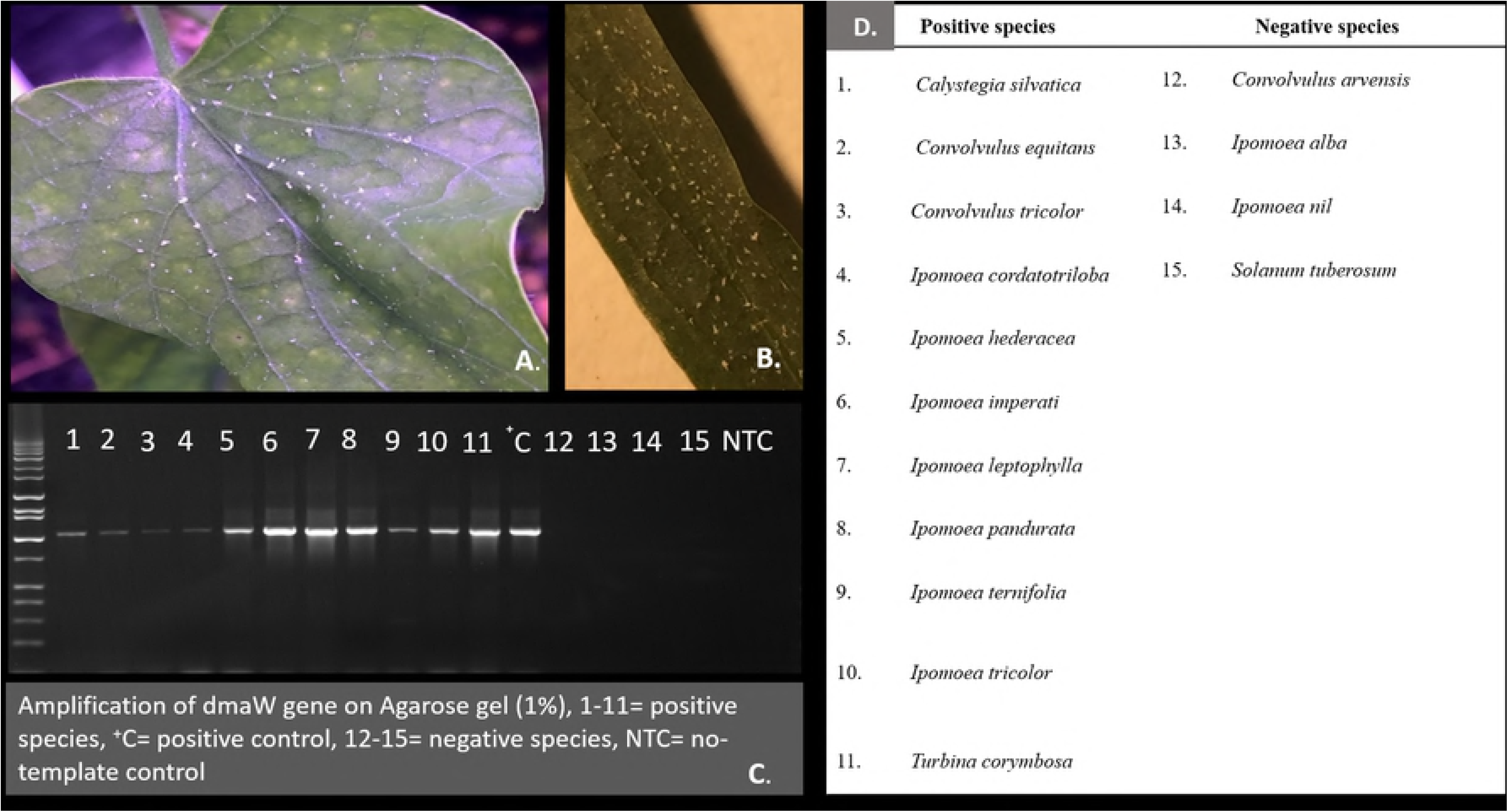
Colonies of *Periglandula* spp. on. (A) *Turbina corymbosa* and (B) *Ipomoea leptophylla*, (C) Agarose gel showing detection of *Periglandula dmaW* gene ~ 1050bp amplicon, (D) List of species in which the *dmaW* gene was detected or not detected corresponding to lane numbers designated in the gel picture.

Ergot alkaloids are categorized into three classes (clavines, simple amides of lysergic acid, and ergopeptines) based on their structural complexity and occurrence in the biochemical pathway [61, 62]. Compounds from two classes (clavines, amides of lysergic acid) were detected in leaf tissues of plant species in which the *dmaW* gene (indicating presence of *Periglandula*) was also detected (Table 3). Compounds included eight clavines and two lysergic acid amides (Table 3). No ergopeptines were detected. Additionally, no ergot alkaloids were detected in species not shown to host *Periglandula* (*C. arvensis*, *I. nil*, *I. alba*). However, the presence of *Periglandula* did not always lead to detection of alkaloids in plant tissues. Alkaloid content may vary with plant age or organ, with higher concentrations typically occurring in seeds and seedlings over vegetative parts [46, 63]. This variation, combined with the possibility that ergot alkaloid concentrations can fall below detection limits, may lead to a failure in confirming presence of ergot alkaloids despite detection of *Periglandula* by molecular methods [55].

**Table 3.** Plant species assayed, psyllid survival (Y/N), and detection of ergot alkaloids by HPLC-MS.

| Plant species* | Survival | Clavines |  |  |  |  |  |  |  | Simple Amides of Lysergic Acid |  | Ergopeptides |  |  |
| --- | --- | --- | --- | --- | --- | --- | --- | --- | --- | --- | --- | --- | --- | --- |
|  |  | Chanoclavine | Lysergic acid (µg/g) | Agroclavine (µg/g) | Lysergol (µg/g) | Festuclavine | Elymoclavine | Elymoclavine fructoside | Dihydrolysergol | Ergonovine (µg/g) | Ergine | Ergotamine | Ergocristine | Ergocornine |
| *C. silvatica* (+) | N | - | - | - | - | - | - | - | - | - | - | - | - | - |
| *C. equitans* (+) | N | - | - | - | - | - | - | - | - | - | - | - | - | - |
| *I. imperati* (+) | N | ++ | 4.4 ± 0.5 | 0.3 ± 0.04 | 0.4 ± 0.05 | ++ | ++ | - | ++ | 5.5 ± 1.0 | ++ | - | - | - |
| *I. leptophylla* (+) | N | ++ | 0.8 ± 0.02 | - | - | + | ++ | - | ++ | 7.1 ± 0.3 | ++ | - | - | - |
| *I. pandurata* (+) | N | + | - | - | - | - | - | - | - | - | - | - | - | - |
| *I. tricolor* (+) | N | ++ | 0.9 ± 0.02 | - | - | + | + | ++ | ++ | 0.3 ± 0.01 | ++ | - | - | - |
| *T. corymbosa* (+) | N | ++ | 3.0 ± 0.03 | - | - | - | + | - | + | 0.7 ± 0.04 | ++ | - | - | - |
| *C. arvensis* | Y | - | - | - | - | - | - | - | - | - | - | - | - | - |
| *C. tricolor* (+) | Y | - | - | - | - | - | - | - | - | - | - | - | - | - |
| *I. alba* | Y | - | - | - | - | - | - | - | - | - | - | - | - | - |
| *I. cordatotriloba* (+) | Y | - | - | - | - | - | - | - | - | - | - | - | - | - |
| *I. hederacea* (+) | Y | - | - | - | - | - | - | - | - | - | - | - | - | - |
| *I. ternifolia* (+) | Y | - | - | - | - | - | - | - | - | - | - | - | - | - |
| *I. nil* | Y | - | - | - | - | - | - | - | - | - | - | - | - | - |
| Potato | Y | - | - | - | - | - | - | - | - | - | - | - | - | - |

\**Periglandula* detected (+)

We observed often striking differences in alkaloid profiles between plant species that allowed psyllid development and species on which the psyllid failed to develop (Table 3). Plants in which clavines and amides of lysergic acid were readily detected were invariably fatal to nymphal psyllids (Table 3). Mortality was quite rapid on these species. Nymphs always died as first instars generally within 24-48 h following egg hatch (Kaur and Horton pers. observation). With two exceptions (*C. silvatica*, *C. equitans*), plant species in which alkaloids were not detected allowed egg-to-adult development (Table 3). Psyllids failed to develop successfully on these two species despite a failure to detect alkaloids and despite detection of *Periglandula* in host tissues (Fig 4C, Table 3). Whether ergot alkaloids were actually present, but not detected, is not known. Lack of survival on *C. equitans* may have been caused in part by the plant’s extreme hairiness, as the pubescence was found to interfere with the ability of psyllids to feed and settle (from visual observations). Psyllids did successfully develop on four other species in which the *dmaW* gene was detected (*C. tricolor*, *I. cordatotriloba*, *I. hederacea*, *I. ternifolia*). However, no ergot alkaloids were detected in leaf tissues from these four species, despite presence of the fungus (Table 3).

## Discussion

This study adds to the list of Convolvulaceae that support egg-to-adult development of potato psyllid, and shows conclusively that the psyllid is able to develop on Convolvulaceae other than the two species (*C. arvensis* and *I. batatas*) previously listed in literature accounts [9, 11, 12]. These additional taxa included an ornamental species of *Convolvulus* (*C. tricolor*) likely of Mediterranean origin [64] and five species of New World *Ipomoea*. The *Ipomoea* comprised a mix of species that are grown as ornamentals (*I. alba*, *I. nil*, *I. hederacea*), and two species (*I. cordatotriloba*, *I. ternifolia*) that are present naturally in regions of Central America, Mexico, and the southwestern U.S. [28, 65, 66, 67]. Previous accounts of association between potato psyllid and Convolvulaceae include rearing trials and field observations. Some care must be taken in interpretation of the field records, as field observations can lead to inflated ideas of true host range [7, 11]. We followed published guidelines [7] in defining psyllid “host plant” as a species that allows egg-to-adult development. A failure to appreciate this distinction has led to confusion about the host range of potato psyllid [11]. We obtained egg-laying and egg hatch (presence of nymphs) on all 14 species of Convolvulaceae that were assayed in this study, but development to the adult stage was limited to seven of these species.

Psyllids of the Central and Northwestern haplotypes were identical with respect to what plant species allowed successful development. The haplotypes did differ in development rates on species allowing development, with psyllids of the Central haplotype developing more rapidly than psyllids of the Northwestern haplotype. Other studies have shown that haplotypes of potato psyllid differ in biological traits, including settling and oviposition behavior [68], development rates [25], body size [24, 25], and composition of endosymbiont communities [27]. The Central haplotype developed more rapidly on cultivated and weedy Solanaceae than psyllids of the Northwestern haplotype [25], which is consistent with our observations. It is likely that differences in development times were partly or largely due to differences between haplotypes in body size. Psyllids of the Northwestern haplotype are conspicuously larger than psyllids of the Central haplotype [24, 25] and it seems likely that the size differences translated into differences in development times between the haplotypes.

We examined whether survival of psyllids on a given plant species could be predicted by location of the species in a phylogenetic tree. The Psylloidea have shown the ability to track phylogenetic diversification of plants within lineages, and host switching or dietary expansion in evolutionary or ecological time by psyllids appear to occur most often between phylogenetically related plants species [8, 31, 32]. One outcome of this sort of phylogenetic tracking is the expectation that dietary breadth for a given psyllid species would likely encompass phylogenetically related plant species rather than a mixture of less-related species. The phylogenetic tree developed from our sequencing work is consistent with trees constructed by earlier phylogenetic work for the Convolvulaceae [60]. Our sequences resolved the fourteen assayed species into two clades which fall respectively into two major tribes [60], the Convolvuleae and Ipomoeeae. The Ipomoeeae was further resolved into two clades [60]: the Argyreiinae, which includes one of our assayed species (*Turbina corymbosa*); and the Astripomoeinae clade, which contains the remaining Ipomoeeae (all species of *Ipomoea*) that were assayed. Our data failed to show that developmental success of psyllids was affected by location of plants in our phylogenetic tree. Plant species that allowed development were represented in both Tribes of Convolvulaceae that were assayed here, as were species that failed to allow development (Fig 1).

Observations in the literature indicate that species of Convolvulaceae may often harbor a class of alkaloids (ergot alkaloids) known in grasses to confer resistance to insect herbivory [43, 45, 61]. These compounds are produced in grasses by fungal species in the family Clavicipitaceae (genus *Epichloё*) which have formed a mutualistic relationship with grasses. A similar mutualistic association between Convolvulaceae and clavicipitaceous fungi in a different genus (*Periglandula*) has been shown to explain the presence of ergot alkaloids in Convolvulaceae [35, 36, 37, 38, 39]. The visual presence of fungal colonies on at least some of our targeted species (Fig 4 AB), combined with extensive literature confirming the presence of ergot alkaloids in Convolvulaceae, prompted us to examine whether psyllid development or lack of development was correlated with the presence or absence of ergot alkaloids.

We detected *Periglandula* in a surprisingly large proportion of assayed plants (11 of 14 species), including in two genera (*Convolvulus, Calystegia*) not previously known to host this fungal symbiont. Previous surveys have suggested that the occurrence of ergot alkaloids (and thus this mutualistic association) was limited to the tribe Ipomoeeae and two clades (Argyreiinae and Astripomoeinae) within this tribe (data based on analyses of 46 species) [34, 42]. It has now been estimated that approximately 50% of Ipomoeeae species, or upwards of 450 species worldwide, could contain ergot alkaloids. These observations understandably have led researchers to focus on the tribe Ipomoeeae in efforts to document presence of ergot alkaloids [34, 42, 37] and it is possible that this focus has led workers to substantially underestimate the taxonomic diversity of Convolvulaceae actually harboring ergot alkaloids. The few reports in the literature suggesting that ergot alkaloids in the Convolvulaceae occur outside of the Ipomoeeae, including in *Calystegia* and *Convolvulus*, have been categorized as “unverified” [34]. Our results are the first to demonstrate that the presence of *Periglandula* in Convolvulaceae does indeed extend outside of Ipomoeeae.

Ergot alkaloids representing two classes (clavines and amides of lysergic acid) were detected in five of our assayed plant species (Table 3). Previous literature accounts summarized in Eich (2008) report these same two classes of alkaloids in four of these five species (failing to list only *I. pandurata*). These same accounts identified many of the same specific compounds that were identified in this study [34, 37]. All species in this study which showed presence of ergot alkaloids were shown (with PCR) to also host *Periglandula*. No ergot alkaloids were detected in the three species in which we failed to detect *Periglandula* (*I. nil*, *I. alba*, *C. arvensis*). This result is consistent with other studies of these three species [34].

We invariably observed 100% mortality of psyllid nymphs on species in which ergot alkaloids were detected (Table 3). Mortality occurred very rapidly following egg hatch, generally within 24-48 h of hatch. Assuming that nymphal mortality was due to the presence of these alkaloids, the next logical question is what mode of action explains our results? Absence of development could have been caused by direct toxicity of the alkaloids or because the compounds deter feeding. At this time, we cannot separate these effects. Insecticidal activity of this class of alkaloids could arise from their capacity to act as agonists or antagonists to neurotransmitter receptors and subsequent malfunctioning of the central nervous system [61]. However, it is also possible that the compounds deterred feeding enough that newly hatched nymphs rapidly desiccated and died. An evaluation of these competing effects will require additional assays, likely including assays that allow measurement of feeding rates (e.g., production of honeydew). Studies in which synthetic analogues of targeted compounds are assayed would also be useful, as use of synthesized compounds would allow insect responses to be examined relative to specific concentrations of compounds or to mixtures of compounds [45, 69].

We failed to detect ergot alkaloids in six species that nonetheless were shown by PCR to harbor *Periglandula* (Table 3). It is unclear if the alkaloids were actually present but were not at detectable levels, if ergot alkaloids were present but were different compounds than targeted by our biochemistry work, or if indeed alkaloids were not present at all. Efforts to detect ergot alkaloids in Convolvulaceae can lead to inconsistent results, even in assays of plant species known from previous studies to harbor the chemicals [34, 55]. These inconsistencies may be the consequence of any of a number of factors, including sensitivity of the analytical approach chosen to look for alkaloids, age of the plant seed or conditions under which the seed was stored, age of the plant, which plant structures are examined, and incorrect taxonomic work leading to mistakes in species identification [34, 70]. Alkaloid levels within a single plant may vary with plant structure. Levels in vegetative tissues, as were targeted here, may be lower than levels in other plant parts, such as seed or newly expanded cotyledons [46, 63]. It may be that analysis and extraction of plant structures other than those that were targeted here (the fully expanded leaf) would have led to detection of ergot alkaloids in those species found to harbor *Periglandula* but in which we failed to detect the chemicals. Potato psyllid successfully completed development on five species in which *Periglandula* was present but in which ergot alkaloids were not detected. If ergot alkaloids do have psyllicidal effects, as suggested by our results in Fig 4 C-D and Table 3, then successful development by psyllids on those five *Periglandula*-positive species from which we failed to detect alkaloids may indicate that alkaloids were indeed not present, or that they were at levels low enough to allow psyllid development and to escape biochemical detection.

Symbiotic association between plants and clavicipitaceous fungi is best known for monocotyledonous plants (Poaceae, Cyperaceae and Junaceae), where (as with Convolvulaceae) the symbioses lead to production of ergot alkaloids [41, 71, 72,]. These associations may lead to any of several benefits for the plant, notably protection against herbivores, but including also nondefense type functions such as enhanced growth rates of the plant or increased ability to withstand drought or other abiotic stresses [73, 74, 75, 76]. Observations that benefits to plants may include multiple types of effects, combined with observations showing that these effects are not always predictable across studies, species, or environments, have led to a large body of literature debating the actual evolutionary processes leading to these associations [76, 77, 78, 79]. Our results provide correlative evidence that presence of ergot alkaloids in Convolvulaceae prevents development of psyllid nymphs, suggesting that the *Periglandula*-Convolvulaceae symbiosis does lead to protection of plants against insect herbivores. Our results also showed, however, that presence of the fungus does not necessarily indicate that psyllids would not survive on the plant host, as species in which *Periglandula* was present but from which alkaloids were not detected did allow egg-to-adult development by psyllids. Future studies will include screening of a larger diversity of Convolvulaceae than assayed here, comprising both *Periglandula*-positive and *Periglandula*-negative species, and we believe that this larger study will shed additional light on the role of this fungal symbiosis in affecting fitness of phloem-feeding insects.

## Acknowledgement

We thank Deb Broers and Jen Stout for technical assistance. We are grateful to Joe Munyaneza and Kylie Swisher for reviewing an earlier draft of this manuscript. Psyllids from Weslaco TX (Central haplotype) were shipped to the ARS facility in Washington State under APHIS permit P526P-17-00366.

## References

1. Munyaneza JE (2012) Zebra chip disease of potato: biology, epidemiology, and management. Am. J. Potato Res. 89: 329–350.

2. Teulon DA, Workman PJ, Thomas KL, Nielsen MC (2009) *Bactericera cockerelli*: incursion, dispersal and current distribution on vegetable crops in New Zealand. N.Z. Plant Prot. 62: 136–144.

3. Western Australia Agriculture and Food (2017) Tomato potato psyllid (TPP). https://www.agric.wa.gov.au/tomato-potato-psyllid-tpp. (Accessed September 2017).

4. Richards BL (1928) A new and destructive disease of the potato in Utah and its relation to the potato psylla. Phytopath. 18: 140–141.

5. Richards BL, Blood H (1933) Psyllid yellows of the potato. J. Agric. Res. 46: 189–216.

6. Capinera JL (2001) Handbook of vegetable pests. Academic Press, New York, NY.

7. Burckhardt D, Ouvrard D, Queiroz D, Percy D (2014) Psyllid host-plants (Hemiptera: Psylloidea): resolving a semantic problem. Fla. Entomol. 97: 242–246.

8. Ouvrard D, Chalise P, Percy DM (2015) Host-plant leaps versus host-plant shuffle: a global survey reveals contrasting patterns in an oligophagous insect group (Hemiptera, Psylloidea). Syst. Biodiv. 13: 434–454.

9. Knowlton GF, Thomas WL (1934) Host plants of the potato psyllid. J. Econ. Entomol. 27: 547.

10. Pletsch DJ (1947) The potato psyllid *Paratrioza cockerelli* (Sulc): its biology and control. Montana Experiment Station, Bulletin 446; 95 pp.

11. Martin NA (2008) Host plants of the potato/tomato psyllid: a cautionary tale. The Weta 35: 12–16.

12. Puketapu A, Roskruge N (2011) The tomato-potato psyllid lifecyle on three traditional Maori food sources. Agron. N.Z. 41: 167–173.

13. Romney VE (1939) Breeding areas of the tomato psyllid, *Paratrioza cockerelli* (Sulc). J. Econ. Entomol. 32: 150–151.

14. Wallis RL (1955) Ecological studies on the potato psyllid as a pest of potatoes. United States Department of Agriculture, Technical Bulletin 1107. 24 pp.

15. Jensen AS, Rondon SI, Murphy AF, Echegaray E, Workneh F, Rashed A, et al. (2012) Overwintering of the potato psyllid in the Northwest on *Solanum dulcamara*. Proceedings of the 12th Annual SCRI Zebra Chip Reporting Session, San Antonio, TX. (http://zebrachipscri.tamu.edu/files/2013/04/2012-Proceedings.pdf)

16. Horton DR, Cooper WR, Munyaneza JE, Swisher KD, Echegaray ER, Murphy AF, et al. (2015) A new problem and old questions: potato psyllid in the Pacific Northwest. Am. Entomol. 61: 234–244.

17. Thinakaran J, Horton DR, Cooper WR, Jensen AS, Wohleb CH, Dahan J, et al. (2017) Association of potato psyllid (*Bactericera cockerelli*; Hemiptera: Triozidae) with *Lycium* spp. (Solanaceae) in potato growing regions of Washington, Idaho, and Oregon. Am. J. Pot. Res. 94: 490–499.

18. Horton DR, Miliczky E, Lewis TM, Cooper WR, Munyaneza JE, Mustafa T, et al. (2017) New geographic records for the Nearctic psyllid *Bactericera maculipennis* (Crawford) with biological notes and descriptions of the egg and fifth-instar nymph (Hemiptera: Psylloidea: Triozidae). Proc. Entomol. Soc. Wa. 119: 191–214.

19. Staples GW, Brummitt R.K. (2007) Convolvulaceae, pp. 108-110. *In* V.H. Heywood, R.K. Brummitt, A. Culham, and O. Seberg (eds.), Flowering Plant Families of the World. Firefly Books, Ontario, Canada.

20. Ouvrard D (2017) Psyl’list – the world Psylloidea database. http://www.hemiptera-databases.com/psyllist. Accessed December 2017.

21. Swisher KD, Munyaneza JE, Crosslin JM (2012) High resolution melting analysis of the cytochrome oxidase I gene identifies three haplotypes of the potato psyllid in the United States. Environ. Entomol. 41: 1019–1028.

22. Swisher KD, Henne DC, Crosslin JM (2014) Identification of a fourth haplotype of the potato psyllid, *Bactericera cockerelli*, in the United States. J. Insect Sci. 14(11): 2014; DOI: 10.1093/jisesa/ieu023.

23. Liu D, Trumble JT (2007) Comparative fitness of invasive and native populations of the potato psyllid (*Bactericera cockerelli*). Entomol. Exp. Appl. 123: 35–42.

24. Horton DR, Miliczky E, Munyaneza JE, Swisher KD, Jensen AS (2014) Absence of photoperiod effects on mating and ovarian maturation by three haplotypes of potato psyllid, *Bactericera cockerelli* (Hemiptera: Triozidae). J. Entomol. Soc. Brit. Col. 111: 1–12.

25. Mustafa T, Horton DR, Swisher KD, Zack RS, Munyaneza JE. (2015a) Effects of host plant on development and body size of three haplotypes of *Bactericera cockerelli* (Hemiptera: Triozidae). Environ. Entomol. 44: 593–600.

26. Mustafa T, Horton DR, Cooper WR, Swisher KD, Zack RS, Munyaneza JE. (2015b) Interhaplotype fertility and effects of host plant on reproductive traits of three haplotypes of *Bactericera cockerelli* (Hemiptera: Triozidae). Environ. Entomol. 44: 300–308.

27. Cooper WR, Swisher KD, Garczynski SF, Mustafa T, Munyaneza JE, Horton DR (2015) *Wolbachia* infection differs among divergent mitochondrial haplotypes of *Bactericera cockerelli* (Hemiptera: Triozidae). Ann. Entomol. Soc. Am. 108: 137–145.

28. Kartesz JT (2011) The biota of North America program (BONAP). *North American plant atlas*. (http://bonap.net/NAPA/Genus/Traditional/County)

29. Swisher KD, Arp AP, Bextine BR, Álvarez EA, Crosslin JM, Munyaneza JE (2013b) Haplotyping the potato psyllid, *Bactericera cockerelli*, in Mexico and Central America. Southwestern Entomol. 38: 201–208.

30. Swisher KD, Munyaneza JE, Crosslin JM (2013a) Temporal and spatial analysis of potato psyllid haplotypes in the United States. Environ. Entomol. 42: 381–393.

31. Percy DM, Page RD, Cronk QC (2004) Plant–insect interactions: double-dating associated insect and plant lineages reveals asynchronous radiations. Syst. Biol. 53: 120–127.

32. Percy DM (2003) Legume-feeding psyllids (Hemiptera, Psylloidea) of the Canary Islands and Madeira. J. Nat. Hist. 37: 397–461

33. Becerra JX (1997) Insects on plants: macroevolutionary chemical trends in host use. Science 276: 253–256.

34. Eich E (2008) Solanaceae and Convolvulaceae: Secondary metabolites: biosynthesis, chemotaxonomy, biological and economic significance. Springer, Berlin.

35. Steiner U, Leibner S, Schardl CL, Leuchtmann A, Leistner E (2011) *Periglandula,* a new fungal genus within the Clavicipitaceae and its association with convolvulaceae. Mycologia 103: 1133–1145.

36. Steiner U, Hellwig S, Ahimsa-Müller MA, Grundmann N, Li SM, Drewke C, Leistner E (2015) The key role of peltate glandular trichomes in symbiota comprising clavicipitaceous fungi of the genus *Periglandula* and their host plants. Toxins 7:1355–1373.

37. Beaulieu WT, Panaccione DG, Ryan KL, Kaonongbua W, Clay K (2015) Phylogenetic and chemotypic diversity of *Periglandula* species in eight new morning glory hosts (Convolvulaceae). Mycologia. 107:667–78.

38. Leistner E, Steiner U (2018) The genus *Periglandula* and its symbiotum with morning glory plants (Convolvulaceae), pp.131–147. Physiology and Genetics: the Mycota (A comprehensive treatise on fungi as experimental systems for basic and applied research), vol. 15. Springer, Cham.

39. Steiner U, Hellwig S, Leistner E (2008) Specificity in the interaction between an epibiotic clavicipitalean fungus and its convolvulaceous host in a fungus/plant symbiotum. Plant Signal Behav. 3: 704–706.

40. Kucht S, Groß J, Hussein Y, Grothe T, Keller U, Basar S, et al. (2004) Elimination of ergoline alkaloids following treatment of *Ipomoea asarifolia* (Convolvulaceae) with fungicides. Planta 219: 619–625.

41. Schardl CL, Panaccione DG, Tudzynski P (2006) Ergot alkaloids—biology and molecular biology. In: Cordell GA, ed. The alkaloids: chemistry and biology. Vol. 63. New York: Academic Press. p 45–86.

42. Eserman LA, Tiley GP, Jarret RL, Leebens-Mack JH, Miller RE (2014) Phylogenetics and diversification of morning glories (tribe Ipomoeeae, Convolvulaceae) based on whole plastome sequences. Am. J. Bot. 101: 92–103.

43. Clay K, Cheplick GP (1989) Effect of ergot alkaloids from fungal endophyte-infected *grasses* on fall armyworm (*Spodoptera frugiperda*). J. Chem. Ecol.15: 169–182.

44. Potter DA, Tyler Stokes J, Redmond CT, Schardl CL, Panaccione DG (2008) Contribution of ergot alkaloids to suppression of a grass-feeding caterpillar assessed with gene knockout endophytes in perennial ryegrass. Entomol. Exp. Appl. 126: 138–147.

45. Shymanovich T, Saari S, Lovin ME, Jarmusch AK, Jarmusch SA, Musso AM, et al. (2015) Alkaloid variation among epichloid endophytes of sleepygrass (*Achnatherum robustum*) and consequences for resistance to insect herbivores. J. Chem. Ecol. 41: 93–104.

46. Amor-Prats D, Harborne JB (1993) Allelochemical effects of ergoline alkaloids from *Ipomoea parasitica* on *Heliothis virescens*. Chemoecology. 4: 55–61.

47. Brummitt RK (1963) A taxonomic revision of the genus *Calystegia*. Ph.D. thesis, University of Liverpool.

48. Brummitt RK (1980) Further names in the genus *Calystegia* (Convolvulaceae). Kew Bull. 35: 327–334.

49. Brummitt RK (2002) *Calystegia silvatica* (Convolvulaceae) in Western North America. Madroño 49: 130–131.

50. Jagoueix S, Bové JM, Garnier M. (1996) PCR detection of the two ‘*Candidatus* Liberibacter species’ associated with greening disease of citrus. Mol. Cell. Probes 10: 43–50.

51. Zhang YP, Uyemoto JK, Kirkpatrick BC (1998) A small-scale procedure for extracting nucleic acids from woody plants infected with various phytopathogens for PCR assay. J. Virol. Methods 71: 45–50.

52. Chen S, Yao H, Han J, Liu C, Song J, Shi L, et al. (2010) Validation of the ITS2 region as a novel DNA barcode for identifying medicinal plant species. PLoS ONE 5: e8613.

53. Yu J, Xue JH, Zhou SL (2011) New universal matK primers for DNA barcoding angiosperms. J. Systematics Evol. 49: 176–181.

54. Kearse M, Moir R, Wilson A, Stones-Havas S, Cheung M, Sturrock S, Buxton S, Cooper A, Markowitz S, Duran C, Thierer T (2012) Geneious Basic: an integrated and extendable desktop software platform for the organization and analysis of sequence data. Bioinformatics 28: 1647–1649.

55. Brown AM (2013) Detection methods and phylogenetic investigation of the morning glory associated fungal symbiont, *Periglandula*. M.S. thesis. Southeastern Louisiana University, Louisiana.

56. Sulyok M, Krska R, Schuhmacher R (2007) A liquid chromatography/tandem mass spectrometric multi-mycotoxin method for the quantification of 87 analytes and its application to semi-quantitative screening of moldy food samples. Anal. Bioanal. Chem. 389: 1505–1523.

57. United States Food and Drug Administration (2001) Guidance for industry bioanalytical method validation [WWW Document]. URL http://www.fda.gov/downloads/Drugs/Guidances/ucm070107.pdf (accessed 7.18.14).

58. SAS Institute (2012) SAS version 9.4. SAS Institute, Cary, NC.

59. Gbur EE, Stroup WW, McCarter KS, Durham S, Young LJ, Christman M, et al. (2012) Generalized linear models, pp. 35-58. Analysis of Generalized Linear Mixed Models in the Agricultural and Natural Resources Sciences. Book News Inc., Portland, OR.

60. Stefanović S, Austin DF, Olmstead RG. (2003) Classification of Convolvulaceae: a phylogenetic approach. Syst. Bot. 28: 791–806.

61. Panaccione DG, Beaulieu WT, Cook D (2014) Bioactive alkaloids in vertically transmitted fungal endophytes. Funct. Ecol. 28: 299–314.

62. Florea S, Panaccione DG, Schardl CL (2017) Ergot alkaloids of the family Clavicipitaceae. Phytopath. 107: 504–518.

63. Beaulieu WT, Panaccione DG, Hazekamp CS, Mckee MC, Ryan KL, Clay K (2013) Differential allocation of seed-borne ergot alkaloids during early ontogeny of morning glories (Convolvulaceae). J. Chem. Ecol. 39: 919–30.

64. Wood JR, Williams BR, Mitchell TC, Carine MA, Harris DJ, Scotland RW (2015) A foundation monograph of *Convolvulus* L. (Convolvulaceae). PhytoKeys 51: 1–282.

65. Austin DF (1990) Annotated checklist of New Mexican Convolvulaceae. Sida 14: 273–286.

66. McDonald JA (1995) Revision of *Ipomoea* section *Leptocallis* (Convolvulaceae). Harv. Pap. Bot. 6: 97–122.

67. Austin DF and Huáman Z (1996) A synopsis of *Ipomoea* (Convolvulaceae) in the Americas. Taxon 45: 3–38.

68. Prager SM, Esquivel I, Trumble JT (2014) Factors influencing host plant choice and larval performance in *Bactericera cockerelli*. PLoS ONE 9: e94047.

69. Bacetty AA, Snook ME, Glenn AE, Noe JP, Hill N, Culbreath A, et al. (2009) Toxicity of endophyte-infected tall fescue alkaloids and grass metabolites on *Pratylenchus scribneri*. Phytopathology. 99:1336–45.

70. Amor-Prats D, Harborne JB (1993b) New sources of ergoline alkaloids within the genus Ipomoea. Biochemical systematics and ecology. 4: 455–61.

71. Clay K, Schardl C (2002) Evolutionary origins and ecological consequences of endophyte symbiosis with grasses. Am. Nat. 160: 99–127.

72. White JF Jr, Bacon CW, Hywel-Jones NL, Spatafora JW (2003) Clavicipitalean fungi, evolutionary biology, chemistry, biocontrol, and cultural impacts. Marcel Dekker, New York.

73. Clay K (1988) Fungal endophytes of grasses: a defensive mutualism between plants and fungi. Ecology 69: 10–16.

74. Malinowski DP, Belesky DP (2000) Adaptations of endophyte-infected cool-season grasses to environmental stresses. Crop Sci. 40: 923–940.

75. Brem D, Leuchtmann A (2002) Intraspecific competition of endophyte infected vs. uninfected plants in two woodland grass species. Oikos. 96: 281–290.

76. Cheplick GP, Faeth SH (2009) Ecology and evolution of the grass-endophyte symbiosis. Oxford University Press. USA.

77. Faeth SH (2002) Are endophytic fungi defensive plant mutualists?. Oikos 98: 25–36.

78. Saikkonen K, Wäli P, Helander M, Faeth SH (2004) Evolution of endophyte–plant symbioses. Trends Plant Sci. 9: 275–280.

79. Clay K (2009) Defensive mutualism and grass endophytes: still valid after all these years. Defensive mutualism in Microbial Symbiosis. Taylor and Francis Publications, pp. 9–20.

